# High-pass noise suppression in the mosquito auditory system

**DOI:** 10.1101/2025.07.07.663513

**Authors:** D.N. Lapshin, D.D. Vorontsov

## Abstract

Mosquitoes detect sound with their antennae, which transmit vibrations to mechanosensory neurons in Johnston’s organ. However, their auditory system is exposed to low-frequency noise from external sources, such as convective and thermal noise, and internal sources, such as flight-induced noise, which could impair sensitivity. High-pass filters (HPFs) may mitigate this issue by suppressing low-frequency interference before it is transformed into neuronal signals.

We investigated HPF mechanisms in *Culex pipiens* mosquitoes by analyzing the phase-frequency characteristics of the primary sensory neurons in the Johnston’s organ. Electrophysiological recordings from male and female mosquitoes revealed phase shifts consistent with high-pass filtering. Initial modeling suggested a single HPF; however, experimental data required revising the model to include two serially connected HPFs to account for phase shifts exceeding –90°.

The results showed that male mosquitoes exhibit stronger low-frequency suppression (∼32 dB at 10 Hz) compared to females (∼21 dB), with some female neurons showing negligible filtering. The estimated delay in signal transmission was ∼7 ms for both sexes. These findings suggest that HPFs enhance noise immunity, particularly in males, whose auditory sensitivity is critical for mating. The diversity in female neuronal tuning may reflect broader auditory functions in addition to mating, such as host detection. This study provides indirect evidence for HPFs in mosquito hearing and highlights sex-specific adaptations in auditory processing. The proposed dual-HPF model improves our understanding of how mosquitoes maintain high auditory sensitivity in noisy environments.

## Introduction

Mosquitoes detect sound using their feathery antennae. When exposed to sound waves, distal segments of the antenna vibrate in proportion to the velocity of air particle displacement (Clements, 1999). These vibrations are transmitted to the Johnston’s organ (JO) located in the pedicel, a modified second segment of the antenna. The JO contains several thousand primary mechanosensory neurons (PSNs) arranged radially (Boo & Richards, 1975), which convert mechanical vibrations into electrical signals.

Similarly to other animals, the auditory perception of mosquitoes is limited by the noise that affects their auditory organs. For a flying mosquito, the sources of noise can be divided into two categories: external, independent of the mosquito’s own activity, and internal, directly related to its behavior. The first group includes convective noise and thermal noise (Goodfriend, 1977; Kazhan et al., 2015, Lapshin, Vorontsov, 2021), both of which should act on the mosquito antenna, an organ sensitive to the air particle velocity, mostly in the low-frequency range. The irregular low-frequency disturbances may be treated as noise, decreasing the signal-to-noise ratio of the mosquito auditory system. The second group includes vibrational noise synchronized with wing beats, and low frequency disturbances associated with flight and maneuvering. The influence of wingbeat vibrations on mosquito hearing has been studied in detail (Jackson & Robert, 2006; Warren et al., 2009; Arthur et al., 2010; Gibson et al., 2010; Lapshin, 2012a, 2012b), the general conclusion being that they increase rather than decrease the sensitivity to certain frequencies on the basis of nonlinear distortions in the JO.

The efficiency of mechanotransduction in the PSNs critically depends on the stability of the operating point, defined as the condition in which receptor sensitivity is maximized (Nadrowski et al., 2004). This is of particular importance for the function of the active amplification mechanism, which involves a complex of dynein-tubulin molecular motors and membrane mechanosensitive ion channels located in the dendrites of the JO PSNs. This active mechanism significantly amplifies sensitivity to very faint signals (Göpfert, Robert, 2001; Nadrowski et al., 2008) (Fig. 1A). If the sensory system deviates too far from its operating point, active amplification becomes either ineffective or entirely disabled. Consequently, the unimpeded transmission of high-amplitude, low-frequency noise from the antenna to the PSNs can hamper the system’s ability to maintain high sensitivity. This suggests that the mosquito auditory system must incorporate mechanisms that reduce the impact of such noise on the PSNs.

**Figure 1.**
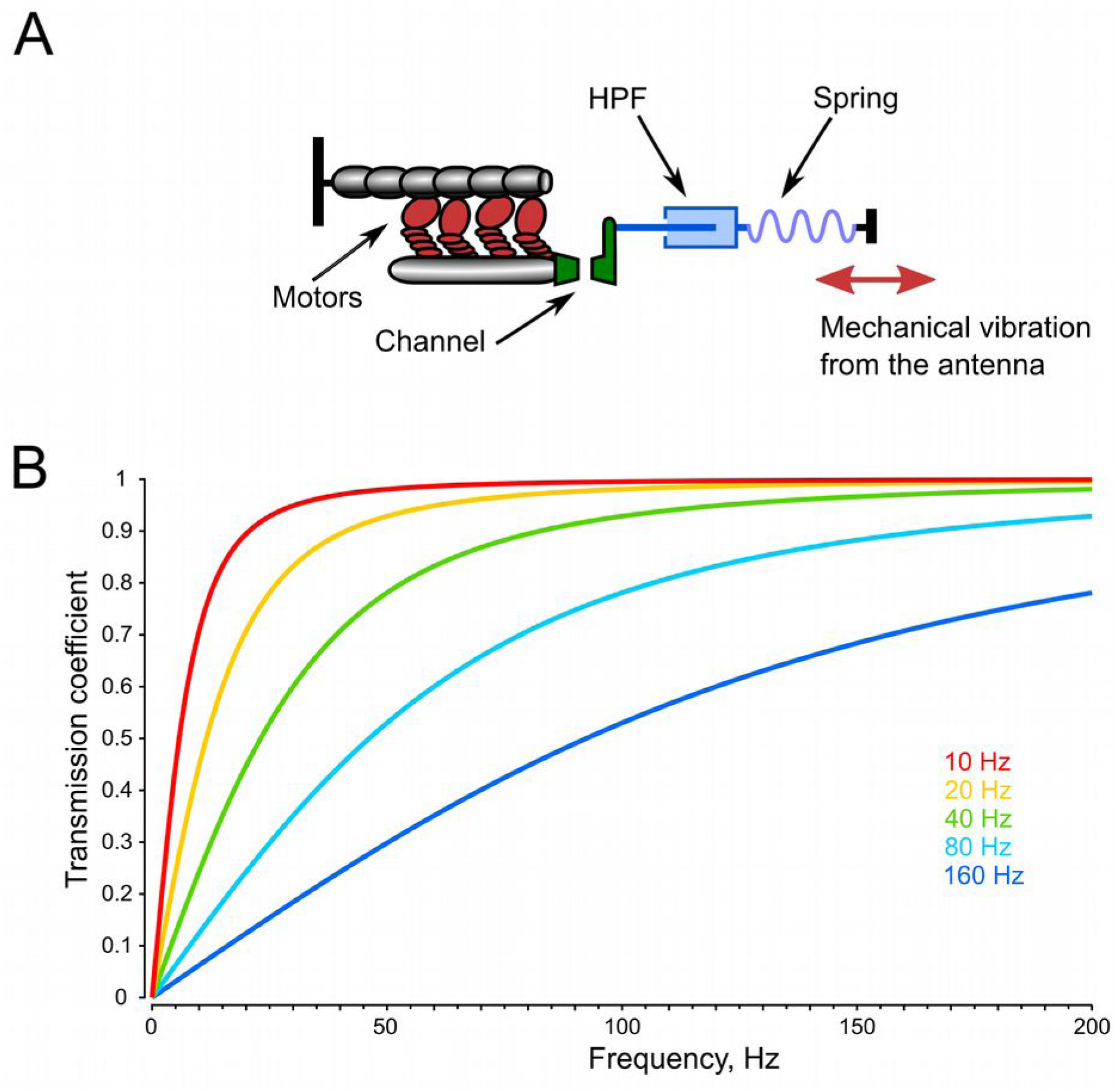
Integration of a high-pass filter into the mechanical transmission pathway from antenna to the active amplification mechanism of the sensory neuron. A – Simplified diagram of the mechanotransduction module, consisting of a single ion channel, an adaptive molecular motor, and an elastic element (adapted from Nadrowski et al., 2008). A high-pass filter (HPF) is added as an additional component in the signal transmission chain. In this model, force is transmitted through a liquid medium; due to viscous properties of liquid, the force at the output of the filter (towards the neuronal membrane with the ion channels) is proportional to the velocity of displacement at the filter input. B – Calculated signal transmission curves of HPFs with different cutoff frequencies from 10 to 160 Hz.

It is well known that many types of sensory receptors act as high-pass filters (HPFs), suppressing stationary or relatively low-frequency signals (Carlson et al., 2019). Such receptors are often referred to as rapidly adapting, velocity-sensitive, dynamic, or phasic mechanoreceptors. A classic example is the Pacinian corpuscle, where rapid adaptation is facilitated by the redistribution of viscous fluid between layers of connective tissue within the corpuscle’s capsule (Bell et al., 1994).

One possible solution to the problem of low-frequency noise suppression is the presence of one or more HPFs (Fig. 1B) along the signal transmission pathway from the antenna to the sensory neurons. These filters can be implemented through mechanical principles or during subsequent electrical signal processing. The design of HPF involves a trade-off between minimizing its influence within the functionally important frequency range and maximizing suppression of out-of-band noise.

While the hypothetic HPFs in the mosquito’s peripheral auditory system cannot be accessed directly, their presence and properties may be inferred indirectly by analyzing the phase shift of electrophysiological responses relative to the acoustic stimuli that elicit them. Under continuous sinusoidal stimulation, the output phase of an HPF leads the input phase in steady-state conditions. Theoretically, the phase shift (i.e., the difference between the output and input phases of a sinusoidal signal) introduced by an HPF increases in magnitude as frequency decreases, approaching –90° at frequencies approaching a few Hertz. As frequency increases, the magnitude of the phase shift gradually decreases, asymptotically approaching zero.

The objective of this study was to identify the signs of high-pass filtering in the peripheral auditory system of mosquitoes and to make preliminary estimates of its physical parameters. To this end, we initially implemented a model comprising a HPF and a delay line arranged in series. A comparison of the modeling results with the experimentally obtained data demanded revising the model by adding a second HPF into the signal chain to achieve consistency with the experimental data.

## Methods

Males and females of *Culex pipiens pipiens* L. were captured in the wild near the Oka river (54° 51’ 44’ N, 38° 21’ 28’ E) in the Moscow region of the Russian Federation.

Electrophysiological experiments were performed under laboratory conditions at an air temperature of 19–21°C at the Kropotovo biological station. A total of 37 experiments were conducted with male and 36 with female mosquitoes.

Each experiment consisted of several sequential stages:

1. Searching for a high-quality recording of the sensory receptor activity in the antennal nerve,
2. Determining the optimal direction of sensitivity in the recording (by rotating the vector of acoustic wave around the axis of the antenna),
3. Measuring the frequency-threshold characteristic (audiogram),
4. Measuring the phase-frequency characteristic.

Individual mosquitoes were fixed by attachment to a 10×5 mm copper-covered triangular plate by a flour paste with addition of sodium chloride, as described in Lapshin, Vorontsov, 2013. Focal extracellular recordings from the axons of the antennal nerve were made with glass microelectrodes (1B100F–4, WPI Inc., Sarasota, FL, USA) filled with 0.15 M sodium chloride and inserted at the scape–pedicel joint. After the penetration of the cuticle, the electrodes had a resistance of 10–60 MΩ.

Neuronal responses were amplified using a home-made AC amplifier (bandpass 5–5000 Hz). For stimulation, two orthogonally oriented stationary speakers were used; they created a vector superposition of acoustic waves at the point of mosquito antenna, as described in detail in Lapshin, Vorontsov (2019). The mosquito was positioned at the crossing of the axes of two speakers in such a way that the antenna’s flagellum was perpendicular to the directions of sound waves originating from each of the two speakers. This approach enabled us to set the desired direction of the acoustic vector relative to the antenna flagellum.

A differential microphone (NR-231-58-000, Knowles Electronics, Itasca, IL, USA) positioned next to the mosquito on a micropositioner with axial rotation feature recorded the stimulation signals. Neuronal responses and stimulation signals were digitized using an Е14-440 A/D board (L-Card, Moscow, Russian Federation) at 20 kHz sampling rate, and LGraph2 software.

Calibration of the stimulating equipment was performed using the same differential microphone. The differential microphone together with its amplifier was previously calibrated in the far field using a B&K 2253 sound level meter with a B&K 4176 microphone (Brüel & Kjær, Nærum, Denmark). All sound level data in this study are given on a logarithmic scale in dB RMS SPVL (root mean square sound particle velocity level), with a reference level of 0 dB being equal to 4.85 × 10⁻⁵ mm/s.

At the beginning of the experiment, as the electrode was gradually advanced into the antennal nerve, the preparation was continuously stimulated with tonal pulses (filling frequency 200 Hz for male mosquitoes, 100 Hz for female mosquitoes, amplitude 60 dB SVPL, duration 80 ms, period 600 ms). In this searching procedure, groups of JO neurons situated orthogonal to the antenna oscillation could be overlooked. To avoid this, the vector of the acoustic wave was periodically changed by 90°.

A response amplitude of 500 µV (peak-to-peak) or more was considered sufficient for subsequent measurements. More detailed methodology for measuring the auditory receptor thresholds was described earlier (Lapshin, 2012a, 2012b; Lapshin, Vorontsov, 2013).

The phasic properties of the auditory response were measured by stimulating the preparation with tonal pulses that incrementally varied from low to high frequencies (50 dB SPVL and 10 Hz increments for male mosquitoes, 60 dB SPVL and 5 Hz increments for female mosquitoes). The duration of individual pulses was typically 3–4 seconds; however, it could be increased in the presence of occasional spontaneous interference in the neuronal response. The interval between successive pulses was maintained at a constant 0.15 seconds.

Both the stimulation control circuit (microphone and the microphone amplifier) and the electrophysiological amplification circuit were pre-calibrated for phase shifts, and the data recorded from the neurons were adjusted accordingly.

## Data Processing

Before the measurement of the phase shift, signals in both recording channels (acoustic stimulation and neuronal response) were bandpass-filtered (see example in Fig. 2) using the Sound Forge 10 PRO software (Sony, Japan). The purpose of this procedure was to isolate the fundamental frequency in each channel while simultaneously suppressing harmonics and noise. The filter was adjusted to the stimulation frequency in each case. To control for artifacts of digital frequency filtering, a pre-synthesized sinusoidal signals of 50 and 100 Hz with predetermined phase shifts between the channels (typically –90°, 90°, and 180°) were filtered in the same way. After the filtering procedure, the phase shift between the signals in the two channels remained unchanged.

**Figure 2.**
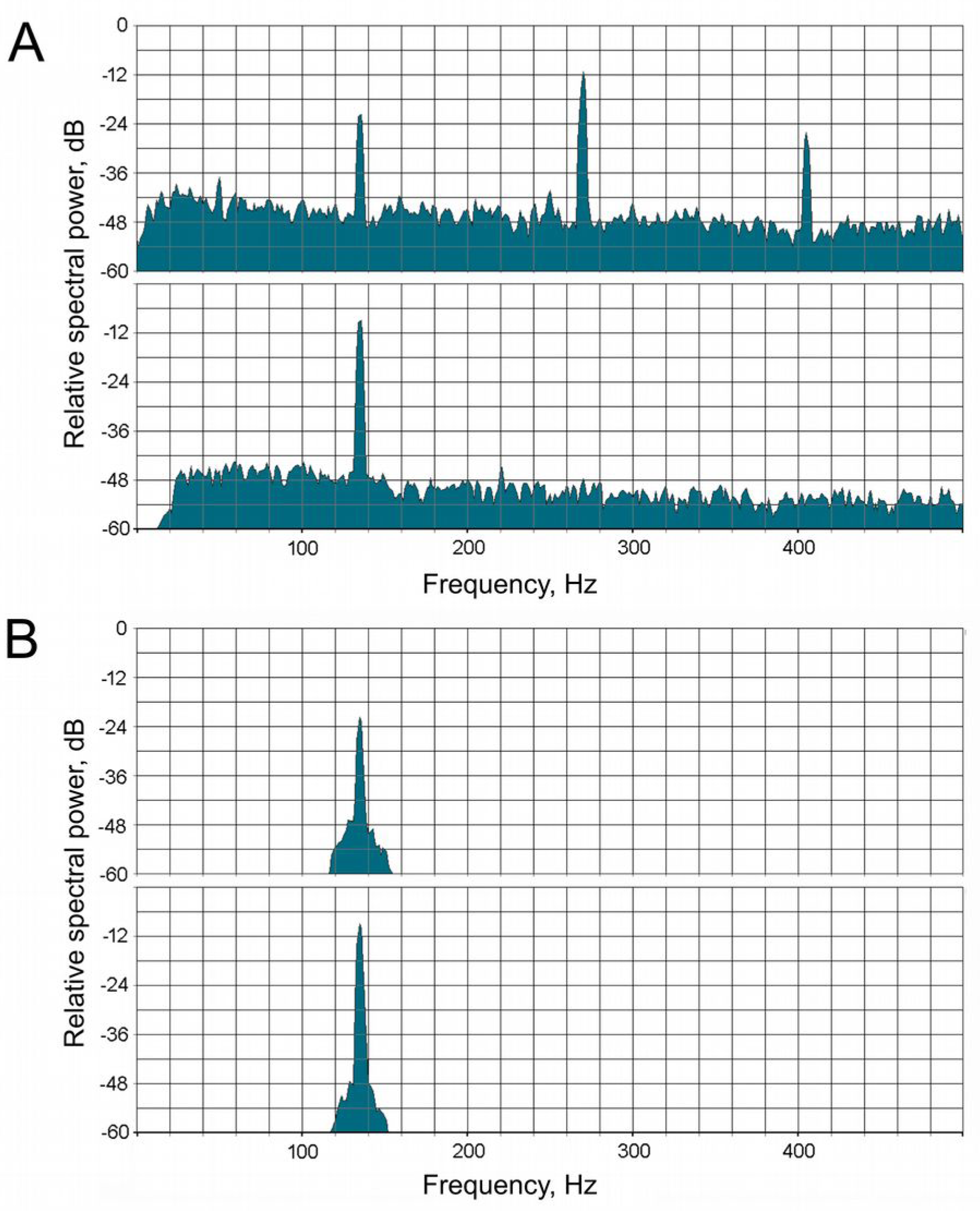
Example of frequency-domain processing of the PSN’s responses in a *Culex pipiens pipiens* female mosquito. A – Spectra of the original electrophysiological response (upper spectrogram) and the acoustic stimulus recorded by the microphone (lower spectrogram). B – the same spectra after digital frequency filtering of the signals, prepared for the phase measurement. The filling frequency of the sinusoidal acoustic stimulus was 135 Hz.

To measure the phase difference between the stimulus and response signals, an instantaneous phase functions for both signals were calculated in Matlab via the Hilbert transform, as was proposed for the analysis of mosquito flight sounds by Aldersley et al. (2014). The phase shift was measured in the second half of a tonal pulse when the system entered a steady-state regime. For each tonal pulse, the median value of the phase shift was taken to plot the phase-frequency characteristic (PFC). The Matlab script and the bandpass-filtered source data are available through Figshare (**please use temporary private link instead of DOI before the article is published in its final form:** https://figshare.com/s/3598059484a7a9faaf5a

10.6084/m9.figshare.29493200).

Linear regression plots were calculated based on the PFCs using the least squares method:

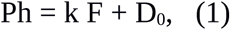

where Ph is the instant phase shift, k is the coefficient characterizing the slope of the regression line relative to the frequency axis, D₀ (initial phase shift) is the point of intersection of regression plot with the vertical axis at F=0.

## Results

### Modeling a system of a high-pass filter and a delay line connected in series

#### High-pass filter

To illustrate the operation of a high-pass filter (HPF), let us consider a system in which oscillations from the antenna are transmitted to the receptor neurons through a fluid medium with a certain viscosity (Fig. 1A). In such a system, the output force of the HPF is proportional to the velocity of displacement of mechanical elements in contact with the fluid. This relationship can be expressed through the following equation:

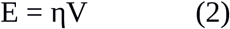

where E is the output force of the HPF, η is a coefficient of viscosity, also including the effects introduced by the geometry and the elasticity of the structures in contact with the fluid, V is the displacement velocity at the input of the filter.

Given that velocity is the first derivative of the displacement function, a system with viscous friction represents a differentiating element. When a sinusoidal signal is applied to the input of such a filter, the output forms the first derivative of a sine function of the same argument, i.e., the product of the angular frequency of the carrier (ω) and a cosine function. Taking into account that ω = 2πF, where F is a carrier frequency, at the time instance t we obtain:

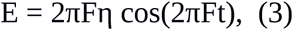

According to this formula, the amplitude of oscillations at the output of the filter is proportional to the frequency of the input signal. At very low frequencies, the transfer coefficient is relatively small, but it increases proportionally with frequency. In real physical systems, the transfer coefficient of an HPF is limited from above, and its general dependence on frequency is shown in Fig. 1B. In other words, an HPF is characterized by a cutoff frequency, below which the signal attenuation increases as the frequency decreases. Above the cutoff frequency, the filter becomes less effective (Fig. 1B). The cutoff frequency is determined, in particular, by the viscosity of the fluid, through which mechanical perturbation is transmitted, and the elasticity of the morphological structures at the input and output of the filter (in our model all these properties are represented by a combined coefficient η).

The second part of the equation (3) — the cosine function—can be viewed as the original sine input function, but phase-shifted by –90°. Theoretically, the phase shift at the HPF output approaches –90° at the lowest frequencies (Fig. 3A).

**Figure 3.**
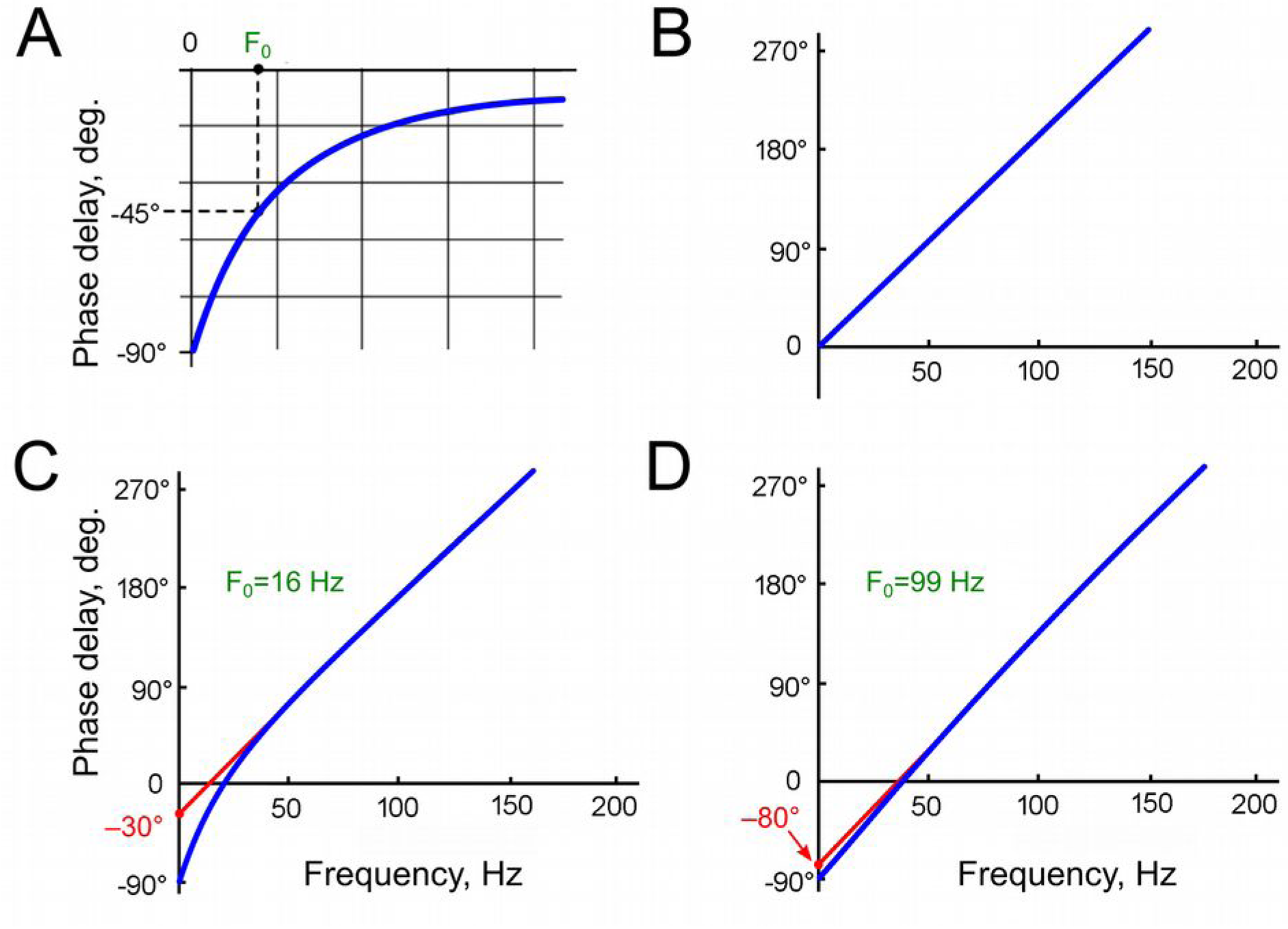
Examples of model phase-frequency characteristics (PFCs, phase shift as a function of frequency). A – General form of the phase-frequency characteristic (PFC) of a first-order high-pass filter (HPF). B – PFC of a delay line (DL, τ₀ = 7.8 ms). C – Resulting phase-frequency characteristic of a HPF with cutoff frequency F₀ = 16 Hz and τ₀ = 7.8 ms. D – Resulting phase-frequency characteristic of a HPF with cutoff frequency F₀ = 99 Hz and τ₀ = 7.8 ms. The regression lines, extrapolated from the phase-frequency plots, intersect the vertical axes at –30° (C) and –80° (D). On experimental plots (Fig. 5), the position of this intersection on the vertical axis (initial phase shift, D₀) reflects the degree of the HPF contribution to the resulting PFC.

#### Delay line

All sensory receptor neurons transfer signals with a certain delay relative to the input (response time delay, τ0). To simplify the reasoning, in our model all mechanisms leading to the signal delay in the axons of receptor neurons relative to the acoustic oscillations affecting the mosquito’s antenna are replaced with a delay line (DL). At the output of the DL, the signal is delayed by time τ0 relative to the input. It is important to note that the DL does not introduce any additional alterations to the signal.

The phase shift at the DL output (Ph2) increases linearly with the frequency of stimulation (Fig. 3B). When the period of the stimulating signal equals the delay time τ0, the total phase shift between the input and output of the DL reaches 360°. From this equality, the response delay τ0 can be determined from the slope of the experimentally measured phase-frequency characteristic (PFC).

At frequencies approaching zero, when the stimulus period is significantly longer than the delay time, the phase shift becomes negligible and approaches zero.

In the model, the parameters that define the DL properties were determined using averaged data on the measured delay time of the electrophysiological response τ, which was calculated from the slope of the experimentally determined PFCs. Given that in a linear system:

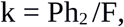

where Ph2 is the phase shift introduced by the DL and F is the signal frequency, we obtain the formula for calculating the delay time from the slope of the experimental PFC:

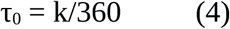

However, it is important to note that the slope of the PFC, and consequently the calculated delay time τ, could be increased by the additional phase shift introduced by the hypothetical HPF. The magnitude of this shift could only be determined after identifying the parameters of this HPF. Consequently, the delay time discussed in relation to the DL refers to the value obtained from the experimental data, unless otherwise stated.

#### Combined phase shift of the model

In a first approximation, we assume that a DL and a hypothetical HPF are connected in series on the path from the antenna to the mechanotransduction area of the receptor neuron. The phase shift at the HPF output (Ph_1_, in angular degrees) is determined by:

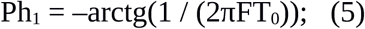

where T_0_ is the time constant of the HPF, F is the signal frequency in Hz. The HPF cutoff frequency F_0_ is related to the time constant T_0_ by:

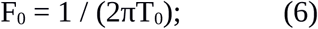

Then:

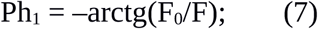

The phase shift introduced by the DL, in angular degrees:

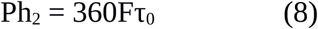

where τ_0_ is the delay in the DL, in seconds. The total phase shift of the system is:

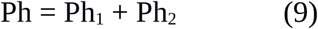

At the lowest frequencies, the phase shift at the HPF output approaches –90°, while the influence of the delay line on the phase characteristic is minimal. In the frequency – phase shift coordinates, the phase characteristic approaches the vertical axis (zero input frequency) at –90° (Fig. 3C,D). At the point where the PFC intersects the vertical axis, the formula (7) does not allow to calculate the angle due to division by zero in the F_0_/F ratio.

As illustrated in Figure 3, the calculated graphs depict two HPFs with arbitrary selected cutoff frequencies of 16 Hz (Fig. 3C) and 99 Hz (Fig. 3D). A comparison of these graphs reveals that, at the higher HPF cutoff frequency, the regression line of the phase characteristic intersects the vertical axis closer to –90°. At frequencies several times higher than the cutoff frequency of the HPF, the DL becomes the predominant factor influencing the total phase shift. Considering the linear dependence of the phase shift introduced by the DL on the frequency, the resulting graph at high frequencies approximates a straight line. If this line is extrapolated toward low frequencies, it should intersect with the vertical axis from –90° to 0°.

The negative value of the phase shift at the point of intersection between the regression line and the vertical axis (i.e., the initial phase shift, D_0_) indicates the presence of the HPF in the system and can be used to estimate the HPF’s cutoff frequency.

### Measurement of phase-frequency characteristics of primary auditory neurons

The experimental testing of the hypothesis regarding high-frequency filtering in the mosquito auditory pathway poses significant methodological challenges. At low frequencies, the sensitivity of the JO receptor neurons undergoes a sharp decrease (i.e., their response thresholds increase). Consequently, the magnitude of the stimulus must be considerably augmented during the measurement of auditory thresholds. However, this approach has limitations: at high amplitudes, nonlinear distortions emerge within the JO, giving rise to harmonic components that are integer multiples of the fundamental stimulus frequency. These components, which appear within the frequency range of the best auditory sensitivity (Fig. 4), elicit neuronal responses to the harmonics rather than to the fundamental frequency of stimulation. This phenomenon imposes constraints on the accuracy of measurements of low-frequency thresholds. Also, at the lowest stimulation frequencies and high amplitudes, analogous nonlinear processes (i.e., the generation of harmonics) emerge in the hardware of the stimulation system, particularly in the acoustic emitters (speakers). In view of the above, there is an area of uncertainty (shaded gray in Fig. 4) within which there are currently no reliable methods for measuring frequency-threshold characteristics within the auditory system of mosquitoes.

**Figure 4.**
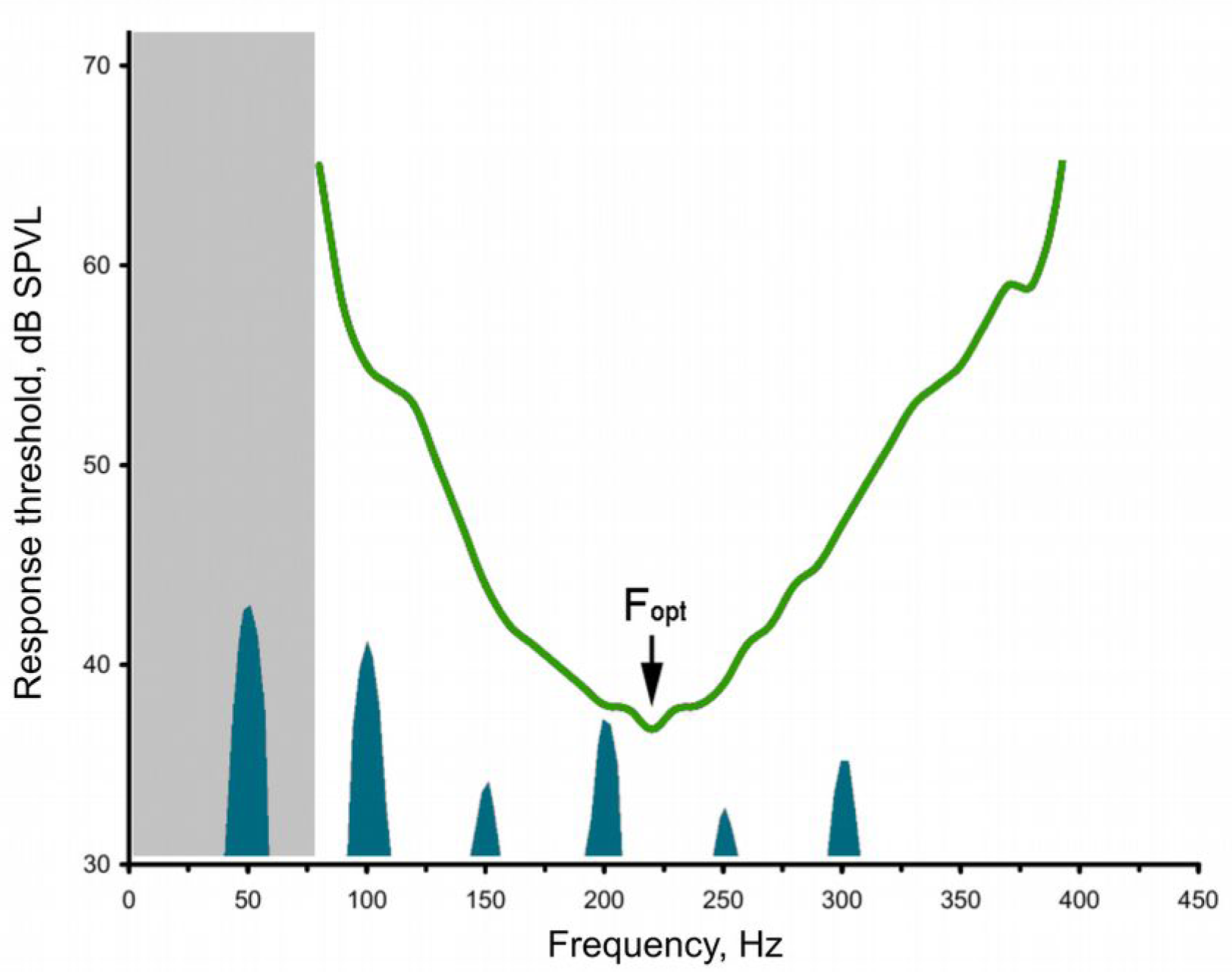
Example of an audiogram of a JO PSN of a male mosquito above the frequency spectrum of harmonic series formed from the 50 Hz sinusoidal signal as a result of nonlinear distortion. Spectral peaks of harmonics are shown below the audiogram curve, some of them fall within the range of maximum auditory sensitivity. The gray-shaded area on the left indicates the frequency range where the auditory thresholds are difficult to measure.

The measurement of phase shifts in the low-frequency range is practically impossible for the same reasons as threshold measurements; namely, the difficulty of obtaining an electrophysiological response that would include the first harmonic of sufficient amplitude to measure the phase difference between the input and output signals. However, using the results of modeling, it became possible to overcome this “uncertainty region” at low frequencies and to integrate the experimental and calculated data into a unified system.

Based on the difference in frequency ranges of male and female mosquitoes (Lapshin, Vorontsov, 2013, 2017), here, in electrophysiological experiments, PFCs were measured in the ranges from 100–160 to 300–360 Hz in male mosquitoes (n=37) and from 50–80 to 180–220 Hz in female mosquitoes (n=36), Marginal segments of the PFC curves that exhibited a substantial deviation from linearity were excluded from further analysis.

It should be noted that the initial phase shift D_0_ can only be determined up to a multiple of 180° (±180·n, where n is an integer). This ambiguity arises because the antennal nerve contains axons from receptor neurons that respond in antiphase to the same antennal deflection (Warren et al., 2010; Lapshin, Vorontsov, 2019, 2023). Figure 5 shows examples of the measured PFCs which are shifted relative to each other by approximately 180°, for pairwise comparison. Data were obtained from males (Fig. 5A) and females (Fig. 5B). Judging by the different slopes of the PFC curves, the receptor neurons differed in their delay times, leading to additional phase shifts under identical stimulation conditions.

**Figure 5.**
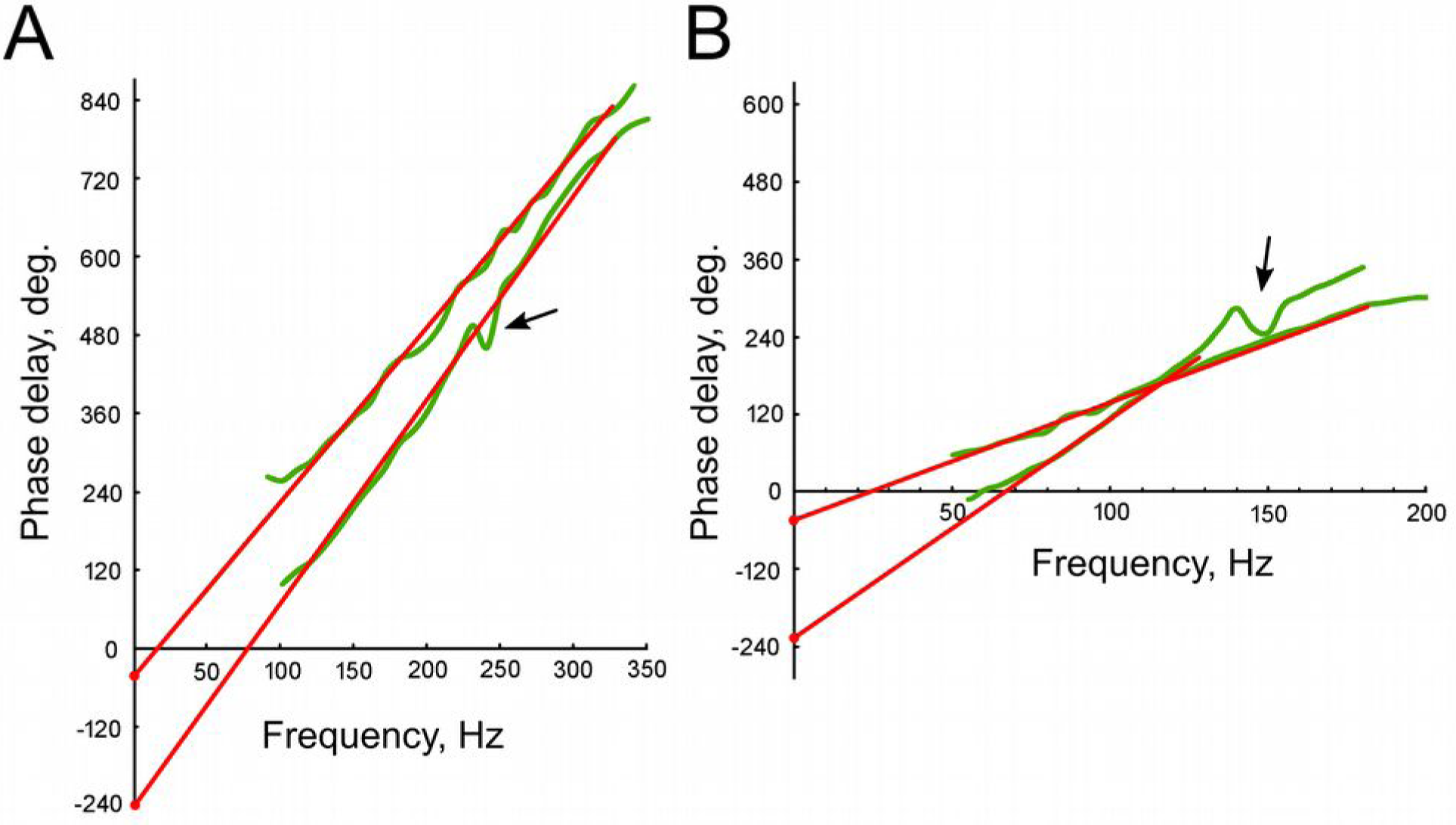
Examples of individual phase-frequency characteristics (PFCs) measured in male (A) and female (B) mosquitoes. The regression plots, extrapolated from the corresponding PFCs, are shown in red. Arrows indicate the regions of increased irregularity of phase shift, which were excluded from subsequent regression analysis. The intersection of the regression plots with the vertical axis reveals the initial phase shift values (D₀).

Wave-like deviations from a relatively straight line were observed in almost all measured PFCs, sometimes reaching significant amplitudes (indicated by arrows in Fig. 5). Such deviations can be attributed to the influence of responses from narrowly tuned receptor neurons on the summary electrophysiological recording (Lapshin, Vorontsov, 2019, 2023). When determining the linear regression parameters, these deviations were excluded. During the subsequent analysis, the values of k (slope) and D_0_ (initial phase shift) were calculated for each experiment, and the delay time τ was then determined from the k values according to the formula (4). The distributions of these parameters is shown in Figure 6, separately for experiments with males (Fig. 6A,C,E,G) and females (Fig.6B,D,F,H).

**Figure 6.**
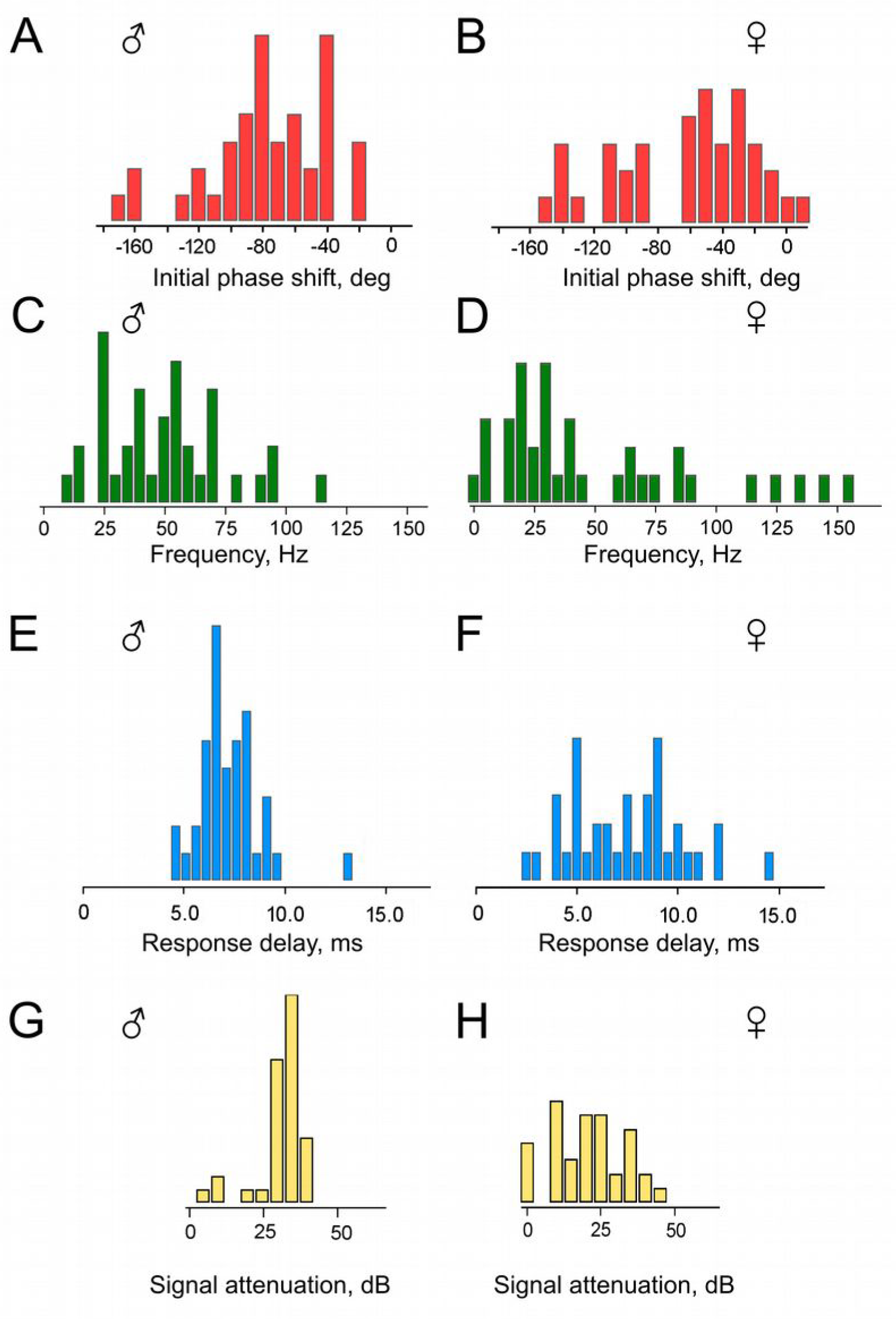
Distributions of parameters calculated on the basis of the experimental data on *Culex pipiens pipiens* mosquitoes. A, B – initial phase shift values, D₀ (bin width of the histograms is 10°). C, D – HPF cutoff frequency values, F₀ (bin width of the histograms is 5 Hz). E, F – delay time values, τ₀ (bin width of the histograms is 1 ms); attenuation levels of a 10 Hz sinusoidal signal at the output of a system composed of two high-pass filters (bin width of the histograms is 5 dB). A, C, E, F – males; B, D, F, H – females.

First, it should be noted that the majority of the obtained D_0_ values fall within the range of negative angles (Fig. 6A,B). This supports our initial hypothesis that the HPF is present in the mosquito auditory system on the way from the antenna to the recording site in the antennal nerve. However, a substantial group of D_0_ values lies below –90°, which is inconsistent with the parameters of a single HPF initially assumed in our calculations (Fig. 3A). The presence of this group of data cannot be explained solely by the scatter of experimental measurements, as each individual D_0_ value was determined from a sequential series of data points forming the PFC, from which the outliers had already been excluded during the preliminary analysis.

The presence of phase shifts D_0_ < –90° can be explained by hypothesizing the existence of at least two HPFs in the signal transmission pathway between the antenna and the auditory receptor axons, potentially based on different physical principles. In such a scenario, the limitation on the maximum negative deviation of the initial phase shift D_0_ should be extended. To test this hypothesis, we re-estimated the total phase shift in the model incorporating two serially connected HPFs.

### Mathematical model of a system including two high-pass filters and a delay line

When two HPFs are connected in series within a functional chain, their transfer coefficients (gains) are multiplied, and the total phase shift is the sum of their individual phase shifts. In this context, it should be noted that the total number of HPFs in the signal chain could potentially exceed two. However, such scenarios cannot be identified using the method employed in this study due to the inherent ±180° uncertainty in estimating the initial phase shift. Thus, here we will consider only one specific case with two HPFs, which have similar cutoff frequencies.

Essentially, another identical HPF is added to the model that already contains a DL and one HPF. Consequently, the maximum phase shift in the low-frequency region doubles, approaching – 180° as F→ 0. The subsequent methodology of analysis remains unchanged.

### Revision of experimental data in view of a model with a DL and two HPFs

Based on the initial values of phase shift (D_0_) and delay time (τ) obtained in each experiment, we calculated: (i) The cutoff frequencies of the two serially connected HPFs (F_0_, Fig. 6C,D), (ii) the delay time τ_0_ calculated according to formula (6) after computationally excluding the phase shift caused by HPFs (Fig. 6E,F).

On average, the calculated delay time τ_0_ was 7.1 ms (SD = 1.5 ms) for males (distribution in Fig. 6E) and 7.4 ms (SD = 2.8 ms) for females (distribution in Fig. 6F). The delay time τ_0_ was 7.5% shorter than the delay time τ measured directly from the slopes of the experimental PFCs. This difference is attributed to the influence of the HPF(s).

To determine the suppression level for a 10 Hz sinusoidal signal, the classic formula for the HPF transfer coefficient (gain) was used:

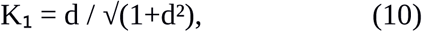

For two serially connected HPFs, their transfer coefficients are multiplied. Given the assumption that the filters have identical parameters:

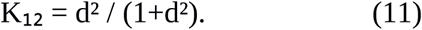

The attenuation level (L) is the reciprocal of the transfer coefficient, expressed in decibels (dB):

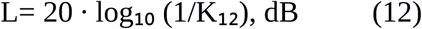

In males, the attenuation level for the 10 Hz signal averaged 32 dB (SD = 5 dB) (Fig. 6G), while in females, it averaged 21 dB (SD = 11 dB) (Fig. 6H).

The data for females includes a group of results with calculated HPF cutoff frequencies clearly below 10 Hz (Fig. 6D); this group corresponds to attenuation levels close to 0 dB (Fig. 6H). In other words, in four experiments with female mosquito’s auditory neurons, they showed virtually no signs of low-frequency filtering.

As an example, we show the attenuation level for a 10 Hz signal at the output of a model system containing two HPFs (Fig. 7).

**Figure 7.**
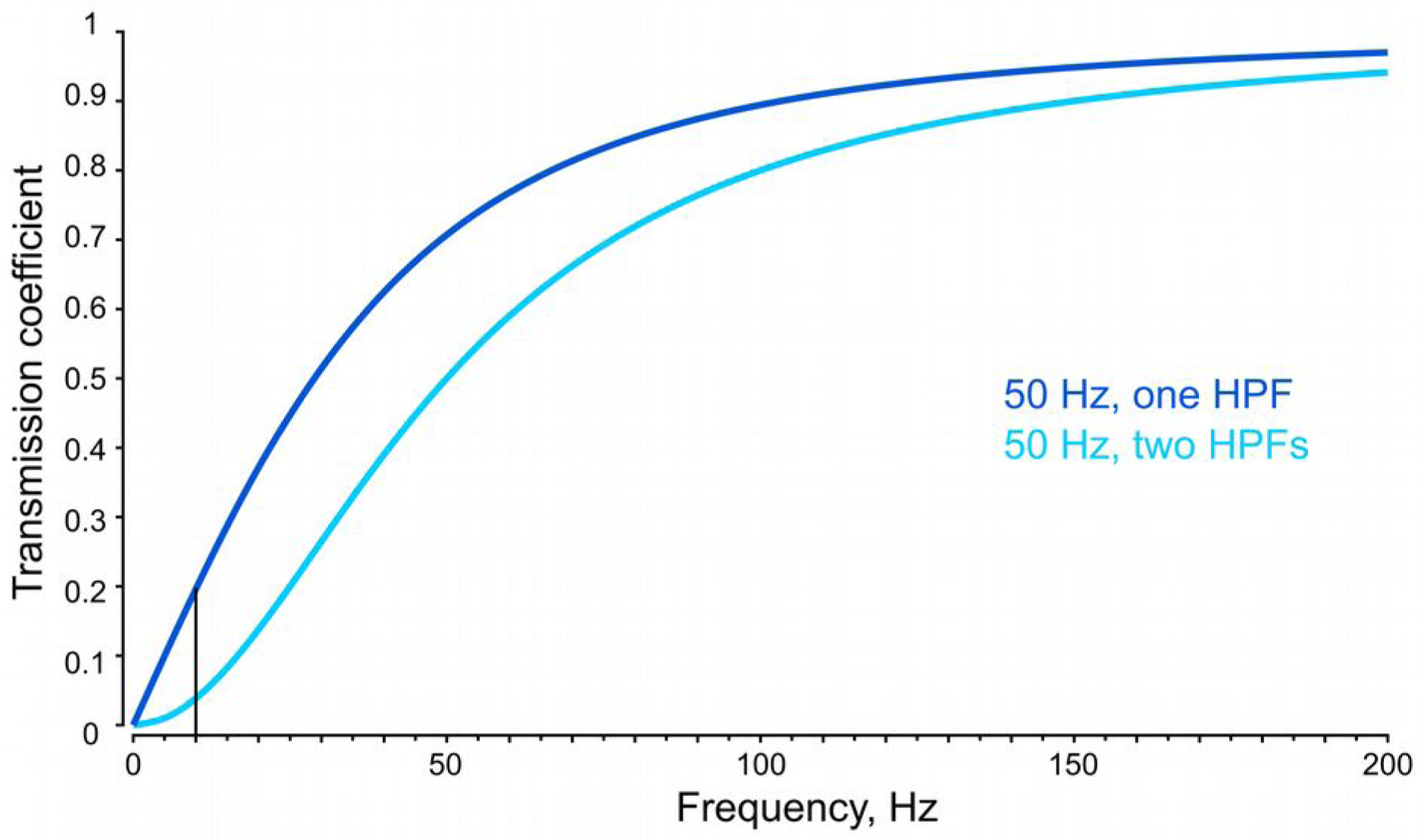
Examples of calculated amplitude-frequency characteristics of a single HPF (A) and two HPFs connected in series (B) with identical cutoff frequency F₀ = 50 Hz, typical for male mosquitoes. Vertical axis shows the transmission coefficient (K) of the system. The level of attenuation at 10 Hz is 14 dB (K₁ = 0.2) for a single HPF and 28 dB (K₁₂ = 0.04) for two HPFs.

## Discussion

According to electrophysiological and behavioral experiments, male mosquitoes possess extremely high auditory sensitivity with response thresholds comparable to those of mammals (Lapshin, Vorontsov, 2021; Feugere et al., 2022). At the same time, such sensitivity is achieved under conditions of high levels of noise from various sources. In particular, at a typical flight speed of 0.4–0.6 m/s during the swarming (Feugère *et al*. 2022) the oncoming uneven airflow that deflects the mosquito’s antennae exceeds the measured auditory thresholds by a factor of a million, or by ca. 120 dB (Lapshin, 2024). It is difficult to imagine a single mechanism capable of ensuring such a high degree of noise immunity. It is more likely that several mechanisms based on different principles are involved at successive stages of perception and processing of acoustic information in the mosquito auditory system, collectively providing the required level of noise filtering.

One such mechanism may be a HPF—or several HPFs—located within the peripheral part of the auditory system. Theoretically expected suppression level (ca. 0.04, or 28 dB at 10 Hz for an HPF with a 50 Hz cutoff frequency, Fig. 7) indicates that such filters suppress a relatively small portion of the low-frequency interference acting on the input of the mosquito auditory system (ca. 120 dB). Nevertheless, their contribution is important, as it reduces the likelihood of overloading the auditory input by the oncoming airflow or gravity on the mosquito’s antennae.

A comparison of attenuation levels in systems with one and with two HPFs showed that in the latter case, the suppression of low-frequency signals was significantly more effective (Fig. 7). Thus, double filtering provides clear advantages in ensuring the noise immunity of the mosquito’s JO. This is especially important for the auditory system of male mosquitoes, whose high auditory sensitivity is crucial for their reproductive success.

High-pass filters do not necessarily have to be incorporated into the mechanical part of the signal processing chain. A similar function may be performed by a complex of ionic mechanisms that stabilize the membrane potential of receptor neurons or other mechanisms responsible for adaptation to slowly changing stimuli.

Our results indicate that characteristics of HPFs are different in male and female mosquitoes. In addition, the auditory system of females includes a group of low-frequency-tuned PSNs without notable signs of high-frequency filtering (Fig.6D,H). These findings are consistent with earlier experiments on females, where no decrease in sensitivity was observed in several PSNs tuned to lowest frequencies (Lapshin, Vorontsov, 2023). However, the frequency characteristics of such neurons can apparently be significantly modified under the influence of neuromodulaion by, for example, octopamine (Vorontsov, Lapshin, 2024).

There is growing evidence that the female auditory system plays a role in both reproductive behavior and the localization of potential hosts by detecting movement or vocalizations (Borkent, Belton, 2006; Bartlett-Healy et al., 2008; Lapshin, Vorontsov, 2023). The capacity to perform such a wide array of tasks has, in turn, given rise to a greater diversity in the characteristics of the JO PSNs. As evidenced by the experimental findings of this study, this diversity manifests in a widened distribution of the HPF cutoff frequencies (Fig. 6C,D), response delay times (latencies) (Fig. 6E,F), and suppression levels of low-frequency signals (Fig. 6G,H) in female mosquitoes compared to males.

In the data obtained from females, the main group of the HPF cutoff frequencies is concentrated in the low-frequency range (Fig. 6D), in contrast to the distribution observed in males (Fig. 6C). This difference is consistent with the data on the neuronal frequency tuning in female (Lapshin, Vorontsov, 2023) and male mosquitoes (Lapshin, Vorontsov, 2019).

Given the relatively wide distribution of individual cutoff frequencies (Fig.6CD), each auditory PSN must contain an HPF located directly at its input. Individual HPFs allow the operating points to be maintained, and thus the high sensitivity, for a large number of sensory neurons despite their different orientations relative to the antennal flagellum. Accordingly, the most likely location of one of the structures implementing high-pass filtering is in the distal part of the dendrite of the auditory neuron within the A1-type sensilla of the JO (Boo, Richards, 1975; Clements, 1999).

## Abbreviations

DL: delay line
JO: Johnston’s organ
HPF: high-pass filter
PFC: phase-frequency characteristic
PSN: primary sensory neuron

## Acknowledgments

The field facilities for this study, Kropotovo biological station, were provided by the Koltzov Institute of Developmental Biology RAS.

## References

Лапшин Д.Н. Слуховая система кровососущих комаров (Diptera, Culicidae). Сенсорные системы, 2024. Т. 38. № 3. С. 3–30. DOI: 10.31857/S0235009224030016

Arthur B.J., Wyttenbach R.A., Harrington L.C., Hoy R.R. Neural responses to one- and two-tone stimuli in the hearing organ of the dengue vector mosquito. *J*. Experimental Biology. 2010. V. 213. P. 1376–1385. DOI: 10.1242/jeb.033357

Aldersley, A.; Champneys, A.; Homer, M.; Robert, D. Time-Frequency Composition of Mosquito Flight Tones Obtained Using Hilbert Spectral Analysis. The Journal of the Acoustical Society of America 2014, 136, 1982–1989, doi:10.1121/1.4895689.

Bartlett-Healy K., Crans W., Gaugler R. 2008. Phonotaxis to amphibian vocalizations in *Culex territans* (Diptera: Culicidae). Annals of the Entomological Society of America, V. 101: 95–103. 10.1603/0013-8746(2008)101[95:PTAVIC]2.0.CO;2

Bell, J.; Bolanowski, S.; Holmes, M.H. The Structure and Function of Pacinian Corpuscles: A Review. Progress in Neurobiology 1994, 42, 79–128, doi:10.1016/0301-0082(94)90022-1.

Borkent A., Belton P. 2006. Attraction of female *Uranotaenia lowii* (Diptera: Culicidae) to frog calls in Costa Rica. The Canadian Entomologist.2006. V. 138: 91–94. 10.4039/n04-113.

Carlson B.A., Sisneros J.A., Arthur N. Popper A.N. Electroreception: fundamental insights from comparative approaches. Springer Handbook of Auditory Research, 2019. 367 p. 10.1007/978-3-030-29105-1

Clements A.N. The biology of mosquitoes Vol. 2 Sensory Reception and Behaviour. New York. CABI Publishing, 1999. 758 p.

Feugère L., Roux O., Gibson G. Behavioural analysis of swarming mosquitoes reveals high hearing sensitivity in *Anopheles coluzzii*. *J*. Experimental Biology. 2022. V. 225. № 5. jeb243535. DOI: 10.1242/jeb.243535

Gibson G., Warren B., Russell I. Humming in tune: sex and species recognition by mosquitoes on the wing. Journal of the Association for Research in Otolaryngology. 2010. V. 11. P. 527–540. DOI: 10.1007/s10162-010-0243-2

Jackson J.C., Robert D. Nonlinear auditory mechanism enhances female sounds for male mosquitoes. PNAS. 2006. V. 103. № 45. P. 16734–16739. DOI: 10.1073/pnas.0606319103

Lapshin D.N. Mosquito bioacoustics: auditory processing in males of *Culex pipiens pipiens* L. (Diptera, Culicidae) during flight simulation. Entomological Review. 2012. V. 92 (6). P. 605–621.

Lapshin D.N. Auditory system of blood-sucking mosquito females (Diptera, Culicidae): acoustic perception during the flight simulation. Entomological Review. 2013. V. 93 (2). P. 135–149.

Lapshin D.N., Vorontsov D.D. Frequency tuning of individual auditory receptors in female mosquitoes (Diptera, Culicidae). J. Insect Physiol. 2013. V. 59. № 8. P. 828–839. DOI: 10.1016/j.jinsphys.2013.05.010

Lapshin D.N., Vorontsov D.D. Frequency organization of the Johnston organ in male mosquitoes (Diptera, Culicidae). J. Experimental Biology. 2017. V. 220. P. 3927–3938. DOI:10.1242/jeb.152017

Lapshin D.N., Vorontsov D.D. Directional and frequency characteristics of auditory neurons in *Culex* male mosquitoes. J. Experimental Biology. 2019. V. 222. jeb208785. DOI: 10.1242/jeb.208785

Lapshin D.N., Vorontsov D.D. Frequency tuning of swarming male mosquitoes (*Aedes communis*, Culicidae) and its neural mechanisms. J. Insect Physiology. 2021. V. 132. DOI: 10.1016/j.jinsphys.2021.104233

Lapshin D.N., Vorontsov D.D. Functions of the auditory system of female mosquitoes (Diptera, Culicidae). Entomological Review, 2023. V. 103: 3, P. 251–262. DOI: 10.1134/S0013873823030016

Lapshin D.N., Vorontsov D.D. Mapping the auditory space of *Culex pipiens* female mosquitoes in 3D. Insects. 2023. V. 14. № 743. P. 1–23. DOI: 10.3390/insects14090743

Nadrowski B., Martin P., Jülicher F. Active hair-bundle motility harnesses noise to operate near an optimum of mechanosensitivity. PNAS. 2004. V. 101. № 33. P. 12195–12200. DOI: 10.1073/pnas.0403020101

Nadrowski B., Albert J.T., Göpfert M.C. Transducer-based force generation explains active process in *Drosophila* hearing. Current Biology. 2008. V. 18. P. 1365–1372. DOI 10.1016/j.cub.2008.07.095

Nadrowski B., Göpfert M.C. Level-dependent auditory tuning: transducer-based active processes in hearing and best-frequency shifts. Communicative and Integrative Biology. 2009. B. 2. Ausgabe 1. S. 7–10. DOI: 10.4161/CIB.2.1.7299

Vorontsov D.D., Lapshin D.N. Effect of octopamine on the frequency tuning of the auditory system in *Culex pipiens pipiens* mosquito (Diptera, Culicidae). Neuroscience and Behavioral Physiology. 2024. V. 54. №. 2. 10 p. DOI 10.1007/s11055-024-01600-2

Warren B., Gibson G., Russell I.J. Sex recognition through midflight mating duets in *Culex* mosquitoes is mediated by acoustic distortion. Current Biology. 2009. V. 9. P. 485–491. DOI: 10.1016/j.cub.2009.01.059

Warren B., Lukashkin A.N., Russell I.J. The dynein–tubulin motor powers active oscillations and amplification in the hearing organ of the mosquito. Proceedings of the Royal Society B. 2010. V. 277. P. 1761–1769. DOI: 10.1098/rspb.2009.2355

